# Integration of haptics and vision in human multisensory grasping

**DOI:** 10.1101/2020.05.12.090647

**Authors:** Ivan Camponogara, Robert Volcic

## Abstract

Grasping actions are directed not only toward objects we see but also toward objects we both see and touch (multisensory grasping). In this latter case, the integration of visual and haptic inputs improves movement performance compared to each sense alone. This performance advantage could be due to the integration of all the redundant positional and size cues or to the integration of only a subset of these cues. Here we selectively provided specific cues to tease apart how these different sensory sources contribute to visuo-haptic multisensory grasping. We demonstrate that the availability of the haptic positional cue together with the visual cues is sufficient to achieve the same grasping performance as when all cues are available. These findings provide strong evidence that the human sensorimotor system relies on non-visual sensory inputs and open new perspectives on their role in supporting vision during both development and adulthood.

## 1 Introduction

The planning and execution of a successful reach-to-grasp movement relies on the translation of visual inputs into motor commands that are involved in the control of the hand (Castiello, 2005; Janssen & Scherberger, 2015; Jeannerod, Arbib, Rizzolatti, & Sakata, 1995). However, actions in everyday life are not only directed toward objects we see but also toward objects we feel with our hands. For instance, even without looking at it, we can easily reach and grasp the cap of a pen while holding the pen with the other hand. Thus, the afferent haptic inputs (proprioception and touch) from the hand in contact with the object are sufficient to estimate the object’s position and its size and successfully guide the action of the opposite hand. However, movements toward haptically sensed objects are usually slower and show a wider grip aperture compared to movements directed toward visually detected objects (Camponogara & Volcic, 2019b; Pettypiece, Culham, & Goodale, 2009; Pettypiece, Goodale, & Culham, 2010). Importantly, the simultaneous availability of vision and haptics leads to faster movements with narrower grip apertures compared to when objects are only visually or haptically sensed (Camponogara & Volcic, 2019a, 2019b). Thus, haptics can effectively integrate with vision to plan and execute grasping actions, but how does each sense contribute to this multisensory-motor transformation is still poorly understood.

Even though vision and haptics provide redundant cues about the position and the size of an object, the two sensory systems acquire and process these cues in fundamentally different ways. In the visual domain, the estimated size of an object is tightly linked to its estimated position in depth. In essence, determining an object’s size requires the scaling of its retinal projections according to its distance from the observer (Brenner & van Damme, 1999; Epstein, Park, & Casey, 1961; van Damme & Brenner, 1997; Volcic & Domini, 2018; Volcic, Fantoni, Caudek, Assad, & Domini, 2013). Instead, in the haptic domain, the estimated size and position of the object are independent. The object size is provided by the proprioceptive and tactile inputs from the digits enclosing the object (Berryman, Yau, & Hsiao, 2006; Durlach et al., 1989; Gaydos, 1958; Langfeld, 1917), whereas the object position is provided by the proprioceptive inputs from the muscles, joints, and skin of the flexed arm (Proske & Gandevia, 2012). Hence, if we assume that the perceived size and position of the objects are used to control grasping movements, there are multiple ways in which visual and haptic cues could be integrated in multisensory grasping. For instance, multisensory grasping could be based on the integration of all visual and haptic position and size cues or only on a subset of these. We captured the different degrees of integration with three models: the full integration, the size integration and the position integration model (Figure 1).

**Figure 1:**
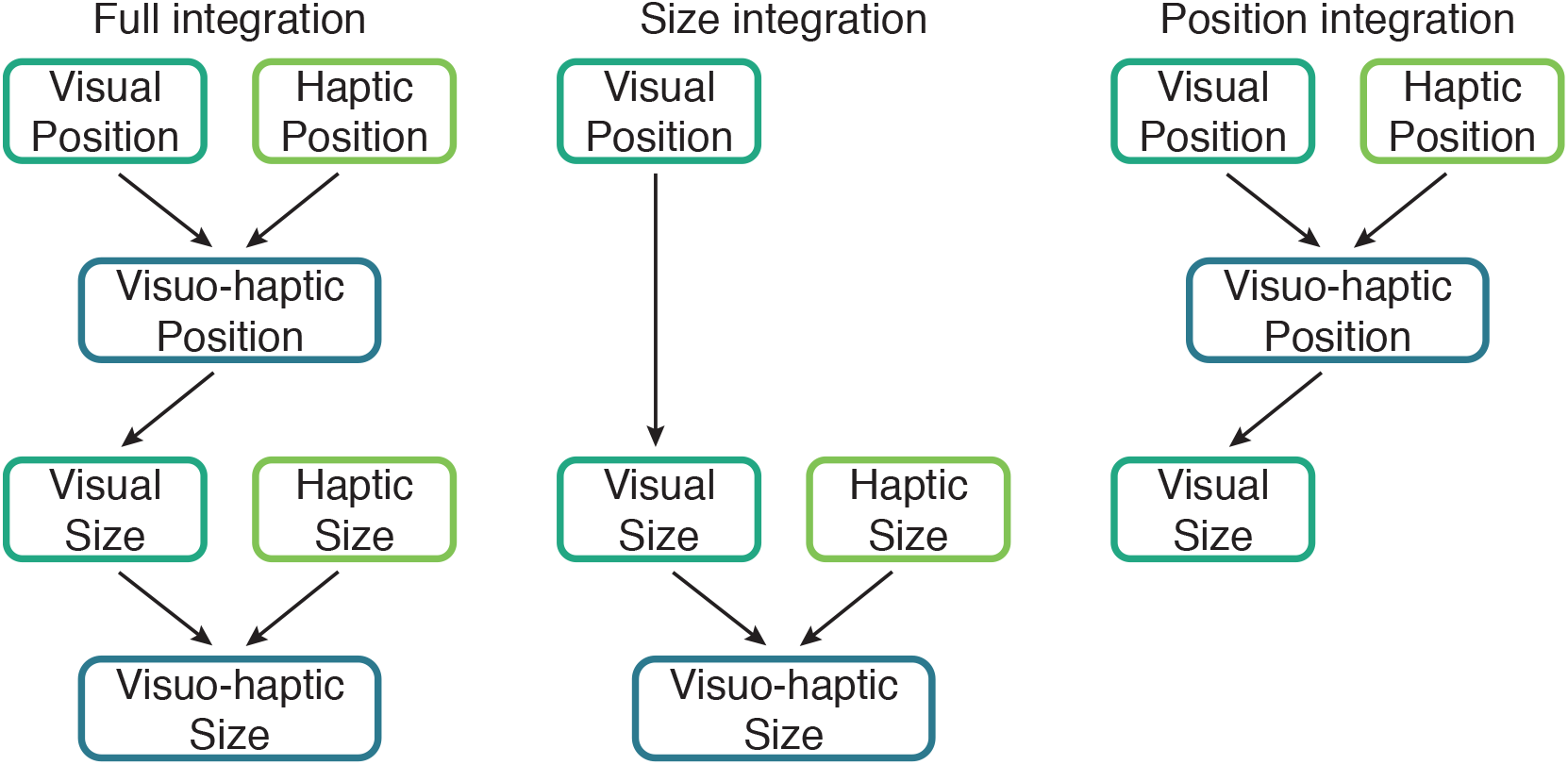
Integration of visual and haptic cues. Left panel, Full integration model: visual and haptic position estimates are merged into a visuo-haptic position estimate. The visual size estimate, which is based on the multisensory position estimate is then integrated with the haptic size estimate. Middle panel, Size integration model: visual object size is mainly based on the visual position estimate. The visual size estimate is then integrated with the haptic size estimate. Right panel, Position integration model: visual and haptic position estimates are integrated into a visuo-haptic position estimate which supports the visual object size estimate.

A *full integration model* posits that both visual and haptic cues about the object size and its position are all merged (Figure 1, left panel): the integration of visual and haptic position estimates promotes a better visual size estimate which is then integrated with the haptic size estimate. The enhanced performance in visuo-haptic multisensory grasping would thus stem from the availability of multisensory estimates of both position and size. An alternative model, the *size integration model* (Figure 1, middle panel), minimizes the role of the haptic position cue and its output is thereby mainly influenced by the integration of haptic and visual size estimates. The joint availability of visual and haptic size would be thus sufficient to improve performance in visuo-haptic grasping. This model resembles the models commonly used to describe visuo-haptic size perception (Ernst & Banks, 2002; Ernst & Bülthoff, 2004; Gepshtein & Banks, 2003; Gepshtein, Burge, Ernst, & Banks, 2005). Another model, the *position integration model* (Figure 1, right panel), is instead focused on the role of haptics as an auxiliary position information cue (Battaglia et al., 2010; Carey & Allan, 1996; Chen, Sperandio, & Goodale, 2018; Sperandio, Kaderali, Chouinard, Frey, & Goodale, 2013, but see Brenner, van Damme, & Smeets, 1997). The output of this model is thus mainly determined by the integration of haptic and visual position estimates with the haptic size cue playing only a marginal role. In this case, the haptic position estimate would improve the visual position estimate which would lead to a better visual size estimate and, in turn, boost visuo-haptic multisensory grasping.

To determine the contribution of haptic and visual cues in multisensory grasping we have characterized grasping performance in two unisensory and two multisensory conditions (Figure 2a). In all conditions the to-be-grasped objects varied in size along the depth dimension and were positioned at different egocentric distances. In the haptic condition (H), vision was prevented, and grasping relied on the haptic size and position cues provided by the left hand holding the object. In the visual condition (V), grasping was based on the visually sensed location and size of the object. In the first multisensory condition, the visuo-haptic condition (VH), both visual and haptic cues about the object position and its size were available. In the second multisensory condition, the visuo-haptic position condition (VHP), participants held with the left hand a post which supported the object, instead of holding the object itself, while vision was fully available. Thus, haptics was informative only about the position of the object, but not about its size. In all these conditions, the grasping action was performed by the right hand.

**Figure 2:**
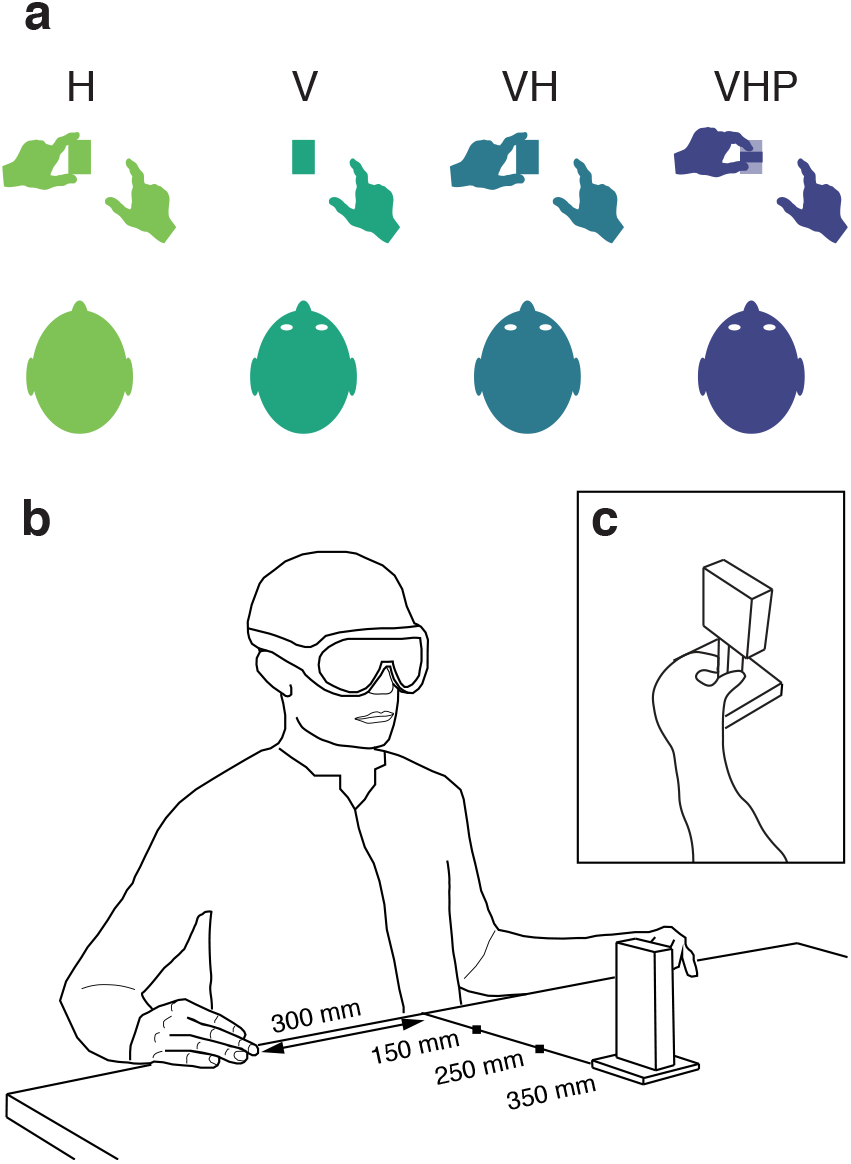
(a) Haptic (H), visual (V), visuo-haptic (VH), and visuo-haptic position (VHP) conditions. Grasping actions were always performed with the right hand. In H, VH and VHP the left hand was already holding the object or the post before the start of the grasping movement. (b) Experimental setup with the stimulus used in conditions H, V, and VH. (c) Left hand holding the post that supported the object in condition VHP.

Based on previous research (Camponogara & Volcic, 2019a, 2019b), we expect the multisensory (VH) condition to exhibit faster grasping movements with smaller peak grip apertures than the V and H unisensory conditions. Whereas all three integration models predict improved grasping performance compared to when objects are only visually or haptically sensed, the comparison between the two multisensory conditions (VH vs. VHP) is instead crucial to distinguish among the full integration, size integration and position integration models. Both the full integration and the size integration models rely, in part, on the availability of the haptic size cue to achieve the multisensory advantage. Thus, if the haptic size cue is absent, both models predict a worse grasping performance in VHP (in which no haptic size cue is provided) than in VH. Grasping movements should be as slow and with peak grip apertures as large as in the V condition according to the size integration model and somewhere in-between the VH and the V conditions according to the full integration model. On the other hand, the position integration model depends mainly on the availability of the haptic position cue. Hence, if only the haptic position cue is sufficient, this model predicts grasping movements in VHP to be as fast and with the same grip aperture as in VH.

## 2 Materials and methods

No part of the study procedures or study analyses were pre-registered prior to the research being conducted. We report how we determined our sample size, all data exclusions, all inclusion/exclusion criteria, whether inclusion/exclusion criteria were established prior to data analysis, all manipulations, and all measures in the study.

### 2.1 Participants

Decisions about the sample size were taken prior to data collection and were based on a recent similar study (Camponogara & Volcic, 2019b) in which the same behavioral paradigm was used. However, we have increased the number of trials per participant to maximize power. Inclusion/exclusion criteria for participants were established prior to the experiment, including normal or corrected-to-normal vision, right-handedness (self-reported), above 18 years of age, and no known history of neurological disorders. No participant was excluded. Twenty students from the New York University Abu Dhabi took part in this study (10 males, age 19.4 ±0.9). All participants were provided with a subsistence allowance. The experiment was undertaken with the understanding and written informed consent of each participant and experimental procedures were approved by the Institutional Review Board of New York University Abu Dhabi.

### 2.2 Apparatus

In the H, V and VH conditions, the stimuli were five 3D-printed rectangular cuboids with a depth of 30, 40, 50, 60, 70 mm, all the same height (120 mm) and width (25 mm) (Figure 2b). In the VHP condition, the stimuli were five 60 mm high rectangular cuboids supported by a 60 mm high post (Figure 2c). While the upper part of these objects varied in depth as in the first set of objects, the depth of the post was kept constant (10 mm) and thus haptics was non-informative about the depth of the to-be-grasped object. In addition, we have manipulated the position of the objects by placing them at three different egocentric distances in front of the participants: at 150 mm, 250 mm or 350 mm in the sagittal direction (depth) at table height. Two 5 mm high rubber bumps with a diameter of 9 mm were attached just in front of the participants, 300 mm to the left and to the right. These bumps were marking the start positions for the left and right hands (Figure 2b).

A pair of occlusion goggles (Red Scientific, Salt Lake City, UT, USA) controlled by a custom Matlab program was used to prevent vision of the workspace in the haptic condition and between trials. A pure tone of 1000 Hz and 100 ms length was used to signal the start of the trial, while another tone of 600 Hz with the same duration was used to signal its end. Index, thumb and wrist movements were acquired on-line at 200 Hz with sub-millimeter resolution by using an Optotrak Certus system (Northern Digital Inc., Waterloo, Ontario, Canada) controlled by the MOTOM toolbox (Derzsi & Volcic, 2018). The position of the tip of each digit was calculated during the calibration phase with respect to three infrared-emitting diodes attached on each distal phalanx (Nicolini, Fantoni, Mancuso, Volcic, & Domini, 2014). An additional marker was attached on the wrist (styloid process of the radius).

### 2.3 Procedure

Participants sat comfortably in front of a table with their torso touching its edge. All the trials started with the participants’ thumb and index digit of the right and left hand positioned on the respective start positions and the shutter goggles closed. Before each trial, one of the objects was positioned at one of the three positions. In the H condition, the experimenter signaled to the participants to hold the object with their left hand along its depth axis at its base (i.e., sense its size and position by means of tactile and proprioceptive inputs) while the shutter goggles remained closed. In the V condition, the goggles turned transparent to provide vision of the object, no haptic information was provided by the left hand. In the VH condition, the participants held the object with their left hand and then the goggles turned transparent. In the VHP condition, participants held the post on which the object was firmly placed with their left hand and then the goggles turned transparent. Unlike in VH, in which haptic inputs were informative about both the object size and its position, the haptic inputs in VHP were informative only about the object’s position (Figure 2c).

After a variable period, the start tone was delivered and participants had to reach for and grasp the object along its depth axis. Movements were performed at a natural speed and no reaction time constrains were imposed. After 3 seconds the end sound was delivered, and, only in the H modality, the goggles were made transparent. Participants had to move their right and left hands back to the start positions and then the goggles turned opaque. Another object and position were then selected and the next trial was ready to start.

The order of conditions was randomized across participants using a Latin square design, while size and position configurations were randomized within each condition. We ran five repetitions for each combination of object size and position, which led to a total of 300 trials per participant (75 for each condition). Before the experiment, a training session was performed in which ten trials were run in each condition to accustom the participants with the task.

### 2.4 Data analysis

Kinematic data were analyzed in R (R Core Team, 2019). The raw data were smoothed and differentiated with a third-order Savitzky-Golay filter with a window size of 21 points. These filtered data were then used to compute velocities and accelerations in three-dimensional space for each digit and the wrist. Movement onset was defined as the moment of the lowest, non-repeating wrist acceleration value prior to the continuously increasing wrist acceleration values (Camponogara & Volcic, 2019b; Volcic & Domini, 2016), while the end of the grasping movement was defined on the basis of the Multiple Sources of Information method (Schot, Brenner, & Smeets, 2010). We used the criteria that the grip aperture is close to the size of the object, that the grip aperture is decreasing, that the second derivative of the grip aperture is positive, and that the velocities of the wrist, thumb and index finger are low. Moreover, the probability of a moment being the end of the movement decreased over time to capture the first instance in which the above criteria were met (Camponogara & Volcic, 2019b; Volcic & Domini, 2016). Trials in which the end of the movement was not captured correctly or in which the missing marker samples could not be reconstructed using interpolation were discarded from further analysis. The exclusion of these trials (90 trials, 1.5% in total) left us with 5910 trials for the final analysis.

We focused our analyses on two dependent variables: the peak grip aperture, defined as the maximum Euclidean distance between the thumb and the index finger, and, the peak velocity of the hand movement, defined as the highest wrist velocity along the movement. These two variables are indicative of the effect of object size and the effect of object position on the movement kinematics (Jeannerod, 1981).

We analyzed the data using Bayesian linear mixed-effects models, estimated using the brms package (Bürkner, 2017) which implements Bayesian multilevel models in R using the probabilistic programming language Stan (Carpenter et al., 2017). The models used to fit the peak grip aperture and peak velocity data included as fixed-effects (predictors) the categorical variable Condition (H, V, VH, VHP) in combination with the continuous variables Size and Position. The continuous variables Size and Position were centered before being entered in the models, thus, the estimates of the Condition parameters (*β_Condition_* intercepts) correspond to the average performance of each Condition. The estimates of the parameters Size (*β_size_* slope) and Position (*β_Position_* slope) correspond instead to the change in the dependent variables as a function of the object size and its position (i.e., distance from the participant). All models included independent random (group-level) effects for subjects. Models were fitted considering weakly informative prior distributions for each parameter to provide information about their plausible scale. We used Gaussian priors for the Condition fixed-effect predictor (peak grip aperture *β_Condition_*: mean = 85 and sd = 40; peak velocity *β_Condition_*: mean = 950 and sd = 500). For the Size and Position fixed-effect predictors we used Cauchy prior distributions centered at 0 with a scale parameter of 2.5. For the group-level standard deviation parameters and sigmas we used Student *t*-distribution priors (peak grip aperture all sd parameters and sigma: *df* = 3, scale = 16; peak velocity sd Condition and sigma: *df* = 3, scale = 166; peak velocity sd Size and sd Position: *df* = 3, scale = 5). Finally, we set a prior over the correlation matrix that assumes that smaller correlations are slightly more likely than larger ones (LKJ prior set to 2).

For each model we ran four Markov chains simultaneously, each for 16,000 iterations (1,000 warm-up samples to tune the MCMC sampler) with the delta parameter set to 0.9 for a total of 60,000 postwarm-up samples. Chain convergence was assessed using the *Ȓ* statistic (all values equal to 1) and visual inspection of the chain traces. Additionally, predictive accuracy of the fitted models was estimated with leave-one-out cross-validation by using the Pareto Smoothed Importance Sampling. All Pareto k values were below 0.5.

The posterior distributions we have obtained represent the probabilities of the parameters conditional on the priors, model and data, and, they represent our belief that the “true” parameter lies within some interval with a given probability. We summarize these posterior distributions by computing the medians and the 95% Highest Density Intervals (HDI). The 95% HDI specifies the interval that includes with a 95% probability the true value of a specific parameter. To evaluate the differences between parameters of two conditions, we have simply subtracted the posterior distributions of *β_Condition_, β_Size_* and *β_Position_* weights between specific conditions (H-V, VH–H, VH–V, VH–VHP). The resulting distributions are denoted as the credible difference distributions and are again summarized by computing the medians and the 95% HDIs.

For statistical inferences about model parameters (*β_Size_* and *β_Position_*) we assessed the overlap of the 95% HDI with zero. A 95% HDI that does not span zero indicates that the predictor has an effect on the dependent variable. For statistical inferences about the differences of the model parameters between conditions, we applied an analogous approach. A 95% HDI of the credible difference distribution that does not span zero is taken as evidence that the model parameters in the two conditions differ from each other.

## 3 Results

We report the main results in two separate sections. In the first section, we describe and compare the performance in the H, V and VH conditions to establish how unisensory and multisensory inputs affects grasping behavior. In the second section, we focus on the critical comparison between the VH and VHP conditions. This comparison allows us to establish whether the haptic size cue is crucial to improve multisensory grasping movements and will reveal which model (full integration, size integration or position integration) is the most plausible.

### 3.1 Grasping with haptic, visual and visuo-haptic inputs

The peak grip aperture was clearly affected not only by the size and position of the object, but, most importantly, also by the available sensory inputs (Figure 3). As can be also seen in Figure 4a, the average peak grip aperture was larger in H (91.8 mm, 95% HDI = 88.2, 95.0) than in V (86.3 mm, 95% HDI = 82.5, 90.3) which, in turn, was larger than in VH (81.2 mm, 95% HDI = 77.7, 84.7). The comparisons between conditions (Figure 4d), showed that the peak grip aperture in H was credibly larger than in both V (H–V = 5.4 mm, 95% HDI = 1.3, 9.4) and VH (H–VH = 10.5 mm, 95% HDI = 6.7, 14.5). Moreover, the peak grip aperture in V was credibly larger than in VH (V–VH = 5.1 mm, 95% HDI = 2.1, 8.0). These peak grip aperture changes were very consistent across participants (Figure 5a, b, and c). The result that the peak grip aperture is smallest in VH replicates and extends previous findings (Camponogara & Volcic, 2019a, 2019b), and supports the idea that the simultaneous availability of visual and haptic inputs leads to a substantial multisensory advantage (approximately 5 mm and 10 mm smaller average peak grip aperture compared to V and H, respectively).

**Figure 3:**
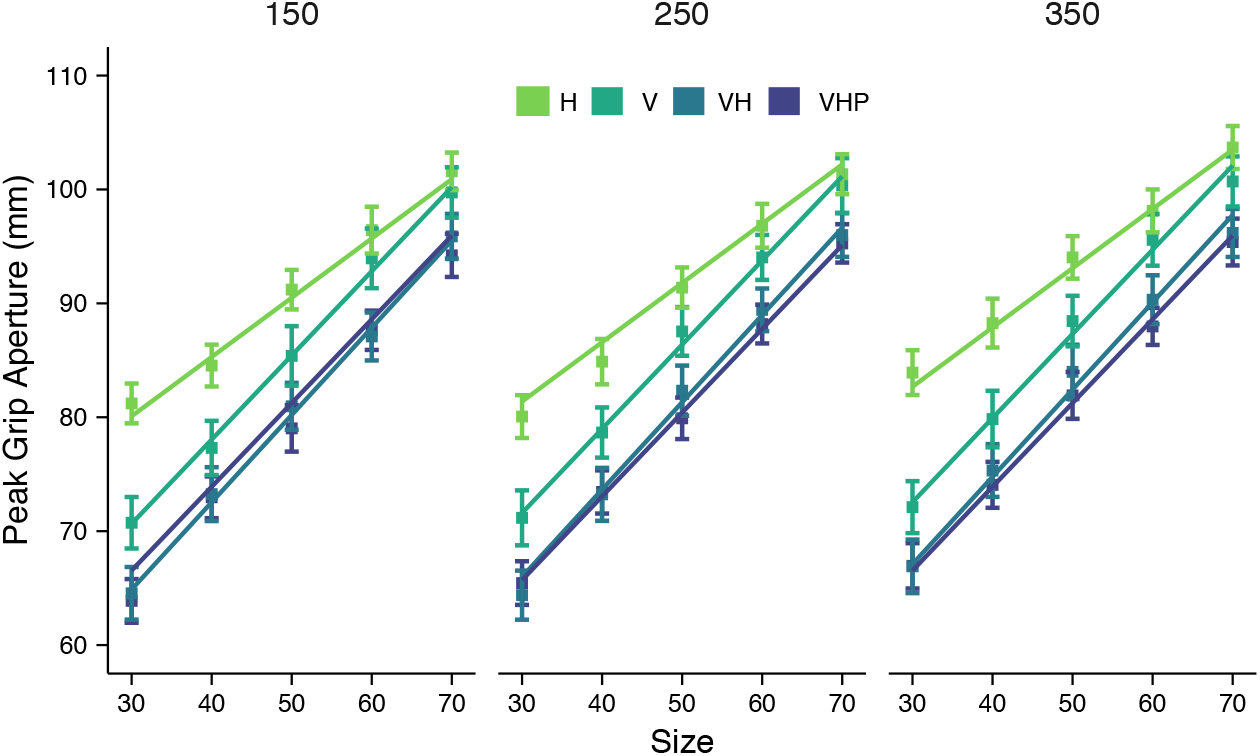
Average peak grip aperture as a function of size and position (separate panels) in the H, V, VH and VHP conditions. Error bars represent the standard error of the mean. Solid lines show the Bayesian mixed-effects model fits.

**Figure 4:**
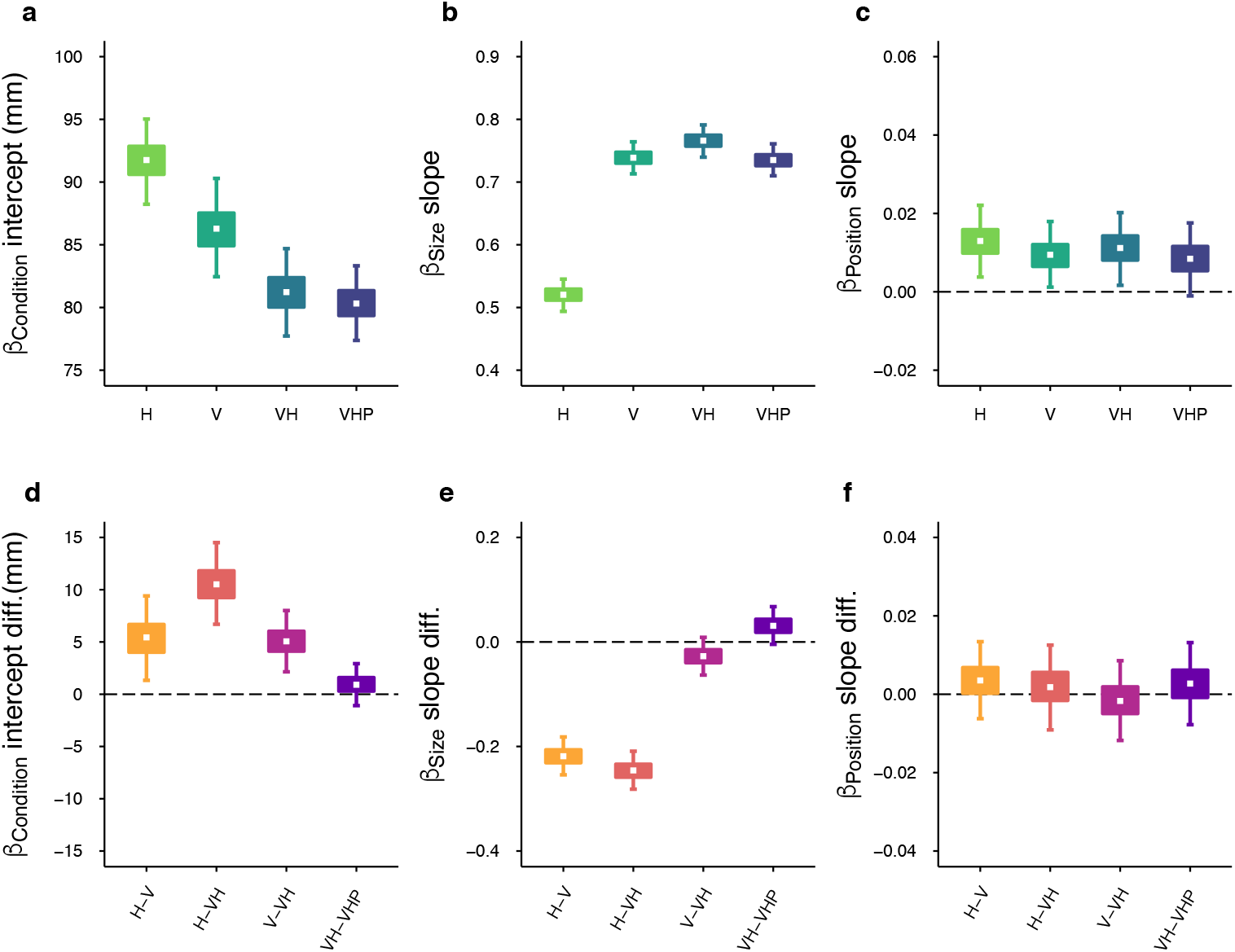
Peak grip aperture results. Top row: Posterior beta weights of the Bayesian linear mixed-effects regression model for the predictors Condition (a), Size (b) and Position (c). Bottom row: Credible difference distributions between conditions for the predictors Condition (d), Size (e) and Position (f). White dots represent the median, the boxes represent the 50% HDIs, and the areas between whiskers represent the 95% HDIs of the distributions.

**Figure 5:**
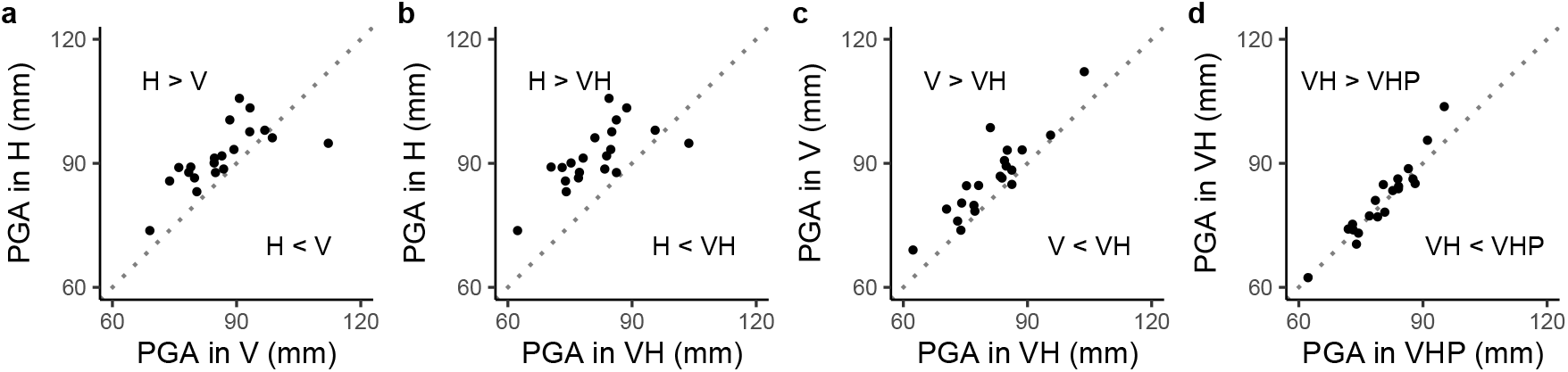
Scatterplots of paired observations. Each point represents the average peak grip aperture (PGA) of a single participant for a pair of conditions: (a) V and H, (b) VH and H, (c) VH and V, (d) VHP and VH. The diagonal reference line of no effect has slope 1 and intercept 0. Points above the diagonal line indicate that the peak grip aperture of the condition represented on the ordinate axis is larger than the peak grip aperture represented on the abscissa.

The peak grip aperture increased as a function of object size in all sensory conditions (Figure 3). However, the slope was shallower in H (0.52, 95% HDI = 0.49, 0.54) than in V (0.73, 95% HDI = 0.71, 0.76) and VH (0.76, 95% HDI = 0.73, 0.80), as can be seen in Figure 4b. The comparisons between conditions (Figure 4e) showed that peak grip aperture increased credibly less as a function of object size in H than in both V (H–V = –0.21, 95% HDI = –0.25, –0.18) and VH (H–VH = –0.24, 95% HDI = –0.28, –0.20). Instead, the changes in peak grip aperture as a function of object size were essentially identical in V and VH (V–VH = –0.02, 95% HDI = –0.06, 0.01). This suggests that the peak grip aperture in H is less sensitive to changes in object size and, thus, the peak grip aperture modulation in multisensory grasping is mainly based on the visual size cue.

The peak grip aperture credibly increased also as a function of object position (Figure 4c), but only moderately (H = 0.012, 95% HDI = 0.003, 0.022; V = 0.009, 95% HDI = 0.001, 0.018, VH = 0.011, 95% HDI = 0.001, 0.02). Most importantly, the comparisons between conditions (Figure 4f) showed that these peak grip aperture increases were indistinguishable between conditions (H–V = 0.003, 95% HDI = –0.005, 0.01; H–VH = 0.001, 95% HDI = –0.008, 0.01; V–VH = –0.001, 95% HDI = –0.01, 0.007).

The peak velocity was mainly affected by the position of the object, but, notably, also by the available sensory inputs (Figure 6). The grasping movements in the unisensory conditions (H = 948 mm/s, 95% HDI = 906, 990; V = 936 mm/s, 95% HDI = 900, 973) reached a lower peak velocity than in the multisensory condition (VH = 971 mm/s, 95% HDI = 948, 1018), as can be also seen in Figure 7a. The comparisons between conditions (Figure 7d) showed that the peak velocity in V was credibly lower than in VH (V–VH = –47 mm/s, 95% HDI = –79, –16). No difference in terms of peak velocity was found between H and V (H–V = 12 mm/s, 95% HDI = –33, 55). Although the credible density distribution of the comparison between H and VH partially overlapped with zero (H–VH = –35 mm/s, 95% HDI = –73, 4), the bulk of the distribution was clearly negative. This pattern of results were very consistent across participants (Figure 8a, b, and c) and suggests that visual and haptic inputs are successfully integrated to speed-up multisensory grasping movements.

**Figure 6:**
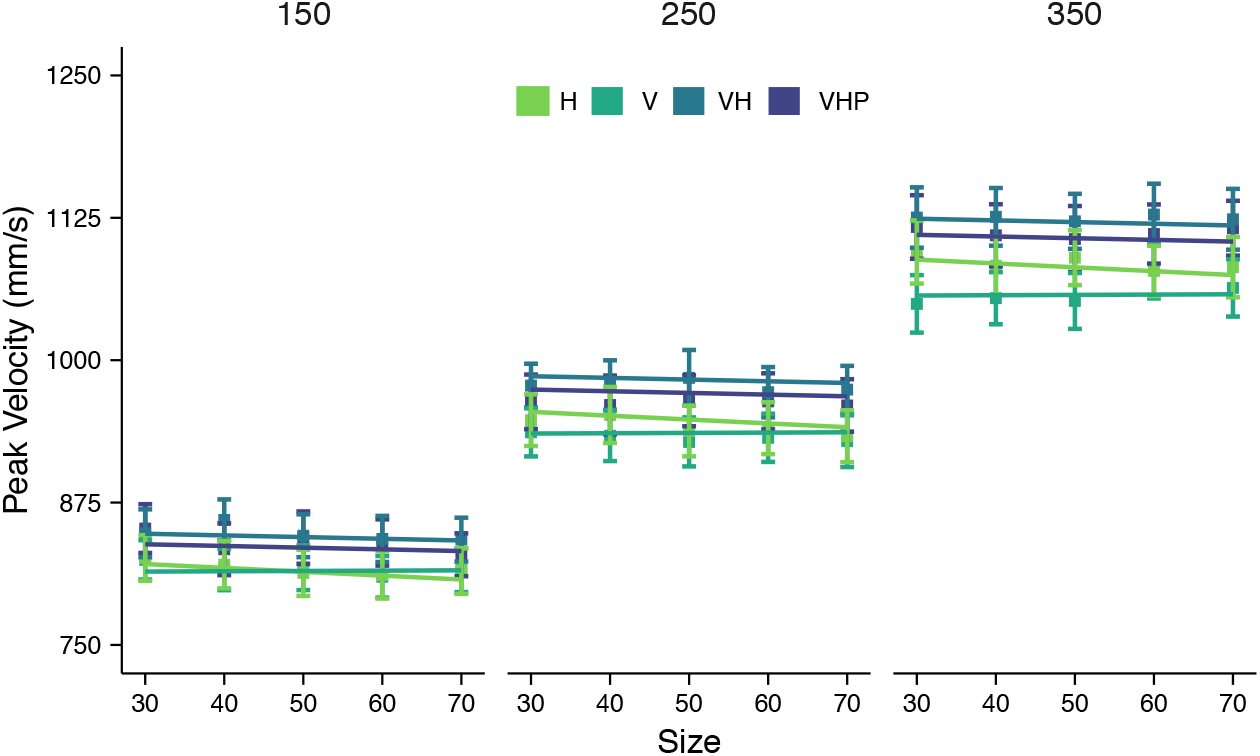
Average peak velocity as a function of size and position (separate panels) in the H, V, VH and VHP conditions. Error bars represent the standard error of the mean. Solid lines show the Bayesian mixed-effects model fits.

**Figure 7:**
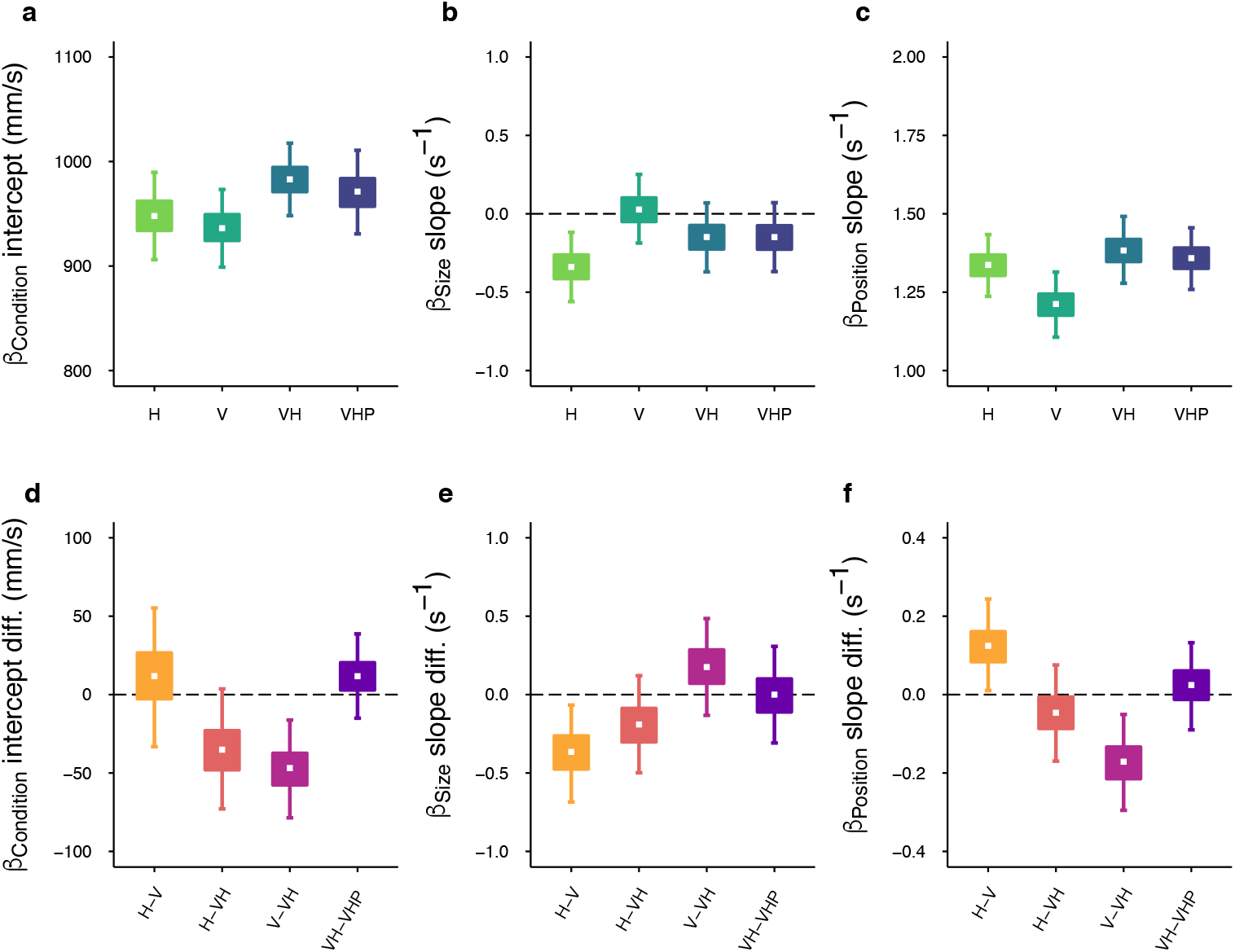
Peak velocity results. Top row: Posterior beta weights of the Bayesian linear mixed-effects regression model for the predictors Condition (a), Size (b) and Position (c). Bottom row: Credible difference distributions between conditions for the predictors Condition (d), Size (e) and Position (f). White dots represent the median, the boxes represent the 50% HDIs, and the areas between whiskers represent the 95% HDIs of the distributions.

**Figure 8:**
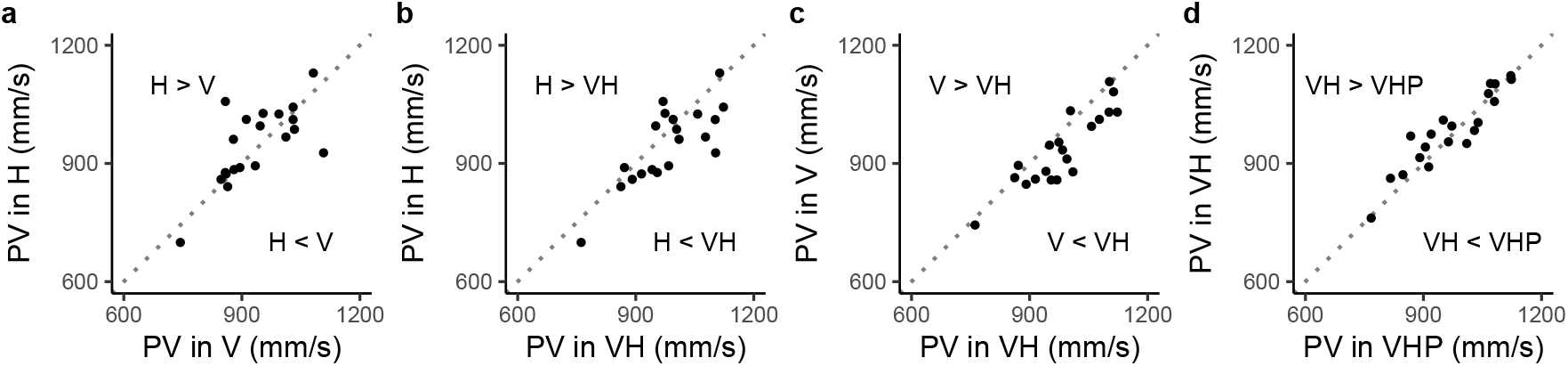
Scatterplots of paired observations. Each point represents the average peak velocity (PV) of a single participant for a pair of conditions: (a) V and H, (b) VH and H, (c) VH and V, (d) VHP and VH. The diagonal reference line of no effect has slope 1 and intercept 0. Points above the diagonal line indicate that the peak velocity of the condition represented on the ordinate axis is larger than the peak velocity represented on the abscissa.

The peak velocity was insensitive to changes in object size in V and VH conditions, as can be gathered from the slopes in Figure 7b, which were relatively flat both in V (V = 0.02 s^-1^, 95% HDI = –0.18, 0.25) and in VH (VH = –0.14 s^-1^, 95% HDI = –0.37, 0.06). Instead, in H, the peak velocity credibly decreased as a function of object size (H = –0.34 s^-1^, 95% HDI = −0.56, –0.12). This decrease might be due to a compensatory strategy specific to haptically-guided grasping wherein participants minimize the risk of colliding with larger objects by decreasing movement velocity instead of increasing the grip aperture. This might be also taken as a symptom of worse sensitivity to size in H than in V. The comparisons between conditions (Figure 7e) showed that the slopes were credibly different only for the comparison between H and V (H–V = –0.36 s^-1^, 95% HDI = −0.68, −0.06). There was no evidence of any other difference (H–VH = −0.19 s^-1^, 95% HDI = –0.49, 0.12; V–VH = 0.17 s^-1^, 95% HDI = –0.13, 0.48).

The object position credibly influenced peak velocity in all sensory conditions (Figure 7c). However, when haptic cues were available, either alone or in combination with vision, the increases in peak velocity as a function of object position were greater than when movements were under visual guidance only (H = 1.33 s^-1^, 95% HDI = 1.23, 1.43; V = 1.21 s^-1^, 95% HDI = 1.10 s^-1^, 1.31; VH = 1.38, 95% HDI = 1.27, 1.49). This result was confirmed by the comparisons between conditions (Figure 7f). Whereas the peak velocity increases in H and VH were credibly different from those found in V (H–V = 0.12 s^-1^, 95% HDI = 0.01, 0.24; V–VH = –0.16 s^-1^, 95% HDI = –0.29, –0.05), no difference was observed between H and VH (H–VH = –0.04 s^-1^, 95% HDI = –0.16, 0.07). These findings provide a first piece of evidence that multisensory integration in grasping relies on the haptic estimates of egocentric object distance more than on the haptic estimates of object size. The comparison between the VH and VHP conditions will further clarify the specific role of the individual haptic cues.

### 3.2 Visuo-haptic grasping with only the haptic position cue

If the haptic size cue is crucial to achieve faster movements with smaller grip apertures in multisensory grasping, then the absence of this cue should deteriorate grasping performance in VHP (Figure 1, left and middle panels) as predicted by the full integration and the size integration models. If, on the other hand, only the haptic position cue is sufficient to boost performance in multisensory grasping, the position integration model predicts that the grip apertures in VHP should be as small as in VH and movements should be as fast (Figure 1, right panel). Indeed, we found that grasping performance in VH and VHP was essentially identical (Figure 3 and 6).

The average peak grip aperture in VHP (VHP = 80.3 mm, 95% HDI = 77.4, 83.3) was the same as in VH (Figure 4d, VH-VHP = 0.9 mm, 95% HDI = −1.1, 2.9) and the peak grip aperture in VHP increased as a function of object size at the same rate as in VH (VHP = 0.73, 95% HDI = 0.70, 0.76), even when the haptic cue about object size was lacking (Figure 4e, VH–VHP = 0.03, 95% HDI = −0.01, 0.06). With regards to object position, the peak grip aperture in VHP increased with distance (0.008, 95% HDI = −0.001, 0.001) as much as in VH (Figure 4f, VH–VHP = 0.002, 95% HDI = −0.006, 0.01). Similarly, the average peak velocity in VHP (971 mm/s, 95% HDI = 931, 1011) was the same as the one we observed in VH (Figure 7d, VH–VHP = 12 mm/s, 95% HDI = −15, 39) and the peak velocity in VHP (VHP = −0.14 s^-1^, 95% HDI = −0.36, 0.07) was as insensitive to changes in object size as in VH (Figure 7e, VH–VHP = −0.0003 s^-1^, 95% HDI = −0.30, 0.30). Finally, in terms of object position, the peak velocity scaling to distance did not differ between VH and VHP (VHP = 1.35 s^-1^, 95% HDI = 1.25, 1.45; VH–VHP = 0.02 s^-1^, 95% HDI = −0.09, 0.13). The similarities between VH and VHP in average peak grip aperture and average peak velocity can be clearly seen also at the per-participant level (Figures 5d and 8d).

The comparable performance in VH and VHP conditions was also visible by tracking the evolution of the grip aperture and wrist velocity along the whole movement (Figure 9a). While the paths of the VH and VHP conditions in wrist velocity-grip aperture space clearly differ from the V and H conditions, they are essentially indistinguishable from each other for all the combinations of object size and position (Figure 9b).

**Figure 9:**
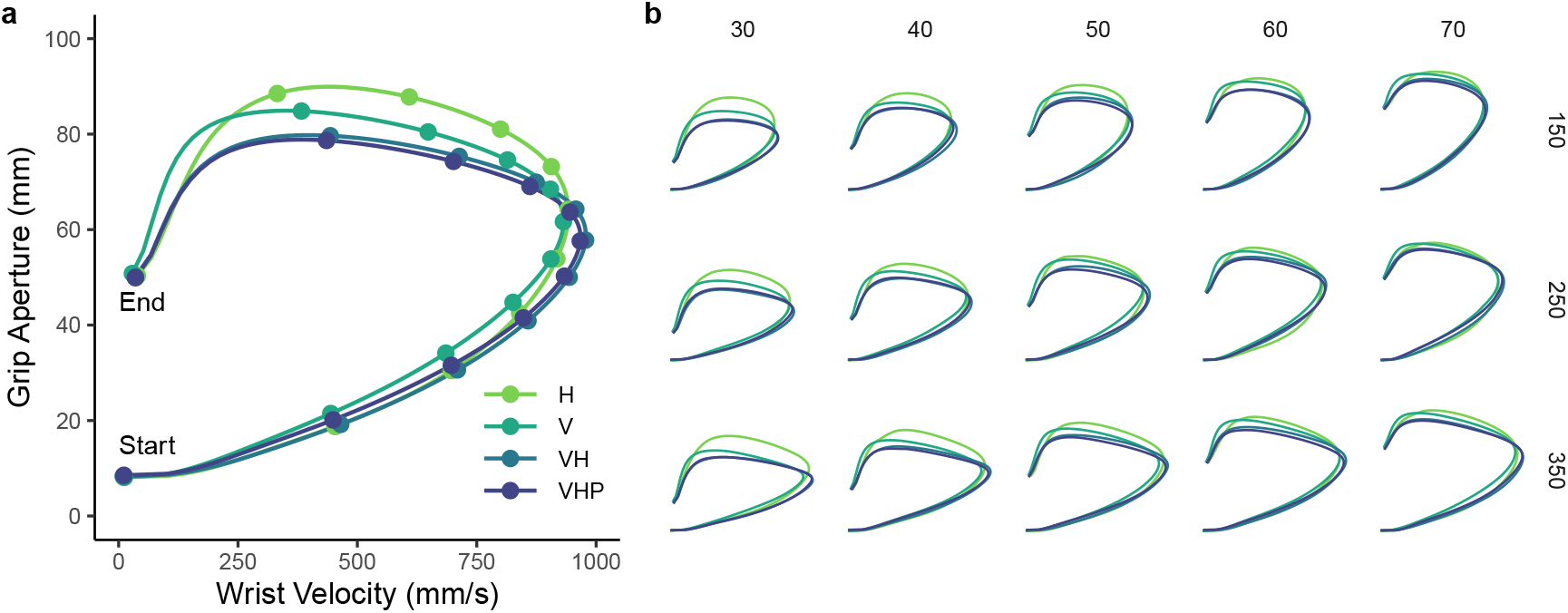
Wrist velocity and grip aperture from start to end of the movement. (a) H, V, VH and VHP conditions are represented by the lines obtained by resampling each movement trajectory in 201 steps evenly spaced along the three-dimensional path and by then averaging the wrist velocity and the grip aperture over all participants, sizes and positions, for each step of the movement trajectory. Points on the lines divide 10% segments equally spaced along the movement trajectory. (b) Same as in (a), but each combination of size and position is represented separately.

Taken together, these results suggest that the haptic position cue, and not haptic size, is required to improve multisensory grasping movements. Thus, faster movements with smaller grip apertures are achieved by integrating visual and haptic position cues that are then jointly used to better scale visual size, consistent with the predictions of the position integration model.

## 4 Discussion

The study presented here was designed to investigate the role of visual and haptic cues in visuo-haptic multisensory grasping. The results show that the availability of both vision and haptics produced faster reach-to-grasp movements with considerably narrower grip apertures than in the unisensory conditions (vision-only or haptics-only) extending our previous findings (Camponogara & Volcic, 2019a, 2019b) over a wide range of object sizes and object positions. Critically, when full vision was coupled with the haptic position cue only (VHP), grasping movements were indistinguishable from those found in the full visuo-haptic condition (VH) in which also the haptic size cue was available.

Because both the full integration and the size integration models rely on the availability of the haptic size cue, its absence should have degraded grasping performance. Our results show that this was evidently not the case, and clearly support the position integration model. Therefore, we must conclude that the main contribution of the haptic modality is related to the estimation of the object position in space. The integration of this haptic position estimate with the available visual information is thus sufficient to enhance grasping performance to the same level as when also the haptic size cue is accessible. The use of haptics in supporting the visual estimation of the object position rather than its size might be related to the intrinsic positional function of the proprioceptive receptors, whose inputs provide a continuous flow of information about the extension of the limbs (Proske & Gandevia, 2012). Our findings thus strengthen the idea that non-visual inputs from the hand holding the object actively support vision in estimating object properties (Battaglia et al., 2010; Carey & Allan, 1996; Chen et al., 2018; Sperandio et al., 2013).

The fact that the haptic size cue might only play a limited role in multisensory grasping might, at first glance, seem surprising. After all, the purpose of multisensory processing is to augment the information contributed by the single modalities (Ernst & Bülthoff, 2004). However, a subordinate role of the haptic size cue could be seen as a virtue rather than a problem, because integrating this specific cue would be in many cases detrimental. Common objects have irregular shapes and, thus, the size perceived by the hand holding the object would rarely match the size of the part of the object that is the target of the grasping action (e.g., reaching with one hand for the cap of a bottle we hold in the other hand). Moreover, actions towards handheld objects do not necessarily end by grasping them (e.g., tapping on a smartphone). In these cases, the haptic position cue can still support vision in movement control, whereas the haptic size cue is essentially irrelevant. By relegating haptic size cues to a secondary role, the sensorimotor system could actually achieve greater robustness to variations in objects shape and action goals.

An additional point that is worth discussing concerns the role of multisensory grasping in learning the mapping between the visual and the motor systems. Infants start to develop the ability to reach and grasp objects at between four and six months of age before learning to perform grasping movements that resemble those of adults at the age of 10–12 months (Gonzalez & Sacrey, 2018; Karl & Whishaw, 2014). Importantly, their inability to produce successful grasps is not due to an immature motor system (Wallace & Whishaw, 2003), to low visual acuity (Banks & Salapatek, 1978) or to undeveloped stereovision (Braddick et al., 1980; Held, Birch, & Gwiazda, 1980), but rather to an unformed mapping between visual inputs and motor plans. This visuomotor mapping is thought to be achieved through an embodied process that requires infants to rely on proprioceptive and, more generally, haptic inputs from their reaching hand (Corbetta, Thurman, Wiener, Guan, & Williams, 2014; Corbetta, Wiener, Thurman, & McMahon, 2018; Thomas, Karl, & Whishaw, 2015). Our findings suggest that valuable haptic spatial cues provided by the hand holding the object could also assist the visuomotor learning process by promoting the development of precise visually controlled reach and grasp movements. In fact, anyone who has witnessed the motor development of a newborn must have observed that they can easily direct a grasping movement toward a handheld object long before they can grasp a distal object they can only see.

It is also important to note that the plasticity of the mapping between visual and motor systems does not end at the onset of adulthood. The visual feedback about the ongoing movement (Bozzacchi, Brenner, Smeets, Volcic, & Domini, 2018; Connolly & Goodale, 1999; Rand, Lemay, Squire, Shimansky, & Stelmach, 2007; Schenk, Mair, & Zihl, 2004; Schettino, Adamovich, & Poizner, 2003; Volcic & Domini, 2016; Winges, Weber, & Santello, 2003), the terminal haptic feedback obtained by the grasping hand (Bingham, Coats, & Mon-Williams, 2007; Bozzacchi, Volcic, & Domini, 2014; Coats, Bingham, & Mon-Williams, 2008; Mon-Williams & Bingham, 2007; Weigelt & Bock, 2007, 2010), and the sensory prediction errors from past trials (Tang, Whitwell, & a. Goodale, 2015; Volcic & Domini, 2018; Whitwell & Goodale, 2009) have all been recognized to be important factors in calibrating visually guided grasping. We suggest that haptic spatial cues about the handheld object could presumably also play a relevant role in maintaining the correct visuomotor calibration over the lifespan or even aid the acquisition of visually guided skills in specific populations, such as children with dense bilateral congenital cataracts who recover vision years after birth (Chen et al., 2016; Held et al., 2011).

Even though this study was not designed to distinguish between different theories about grasping, it provides some important insights to be considered. First, according to the two-visual-systems hypothesis (Goodale, 2011), binocular vision provides estimates of distance and size that are accurate and reliable and is thus critical for grasping guidance. Indeed, switching from binocular to monocular vision has clear detrimental effects on grasping (Bradshaw et al., 2004; Melmoth & Grant, 2006; Servos, Goodale, & Jakobson, 1992). Thus, one might expect that providing a haptic position cue should compensate for the loss of visual distance cues in conditions of limited vision. This is, in fact, what has been found (Chen et al., 2018). Our finding that the haptic position cue reduces the grip aperture and increases movement velocity even in conditions of full binocular vision instead suggests that vision-for-action is less efficient than it is usually claimed. Second, our finding that holding the pole on which the object is placed (i.e., only haptic position cue available) leads to identical grasping performance as when both haptic size and position cues are available is not fully compatible with the theories about grasping that propose that either a single digit (Galea, Castiello, & Dalwood, 2001; Haggard & Wing, 1997; Melmoth & Grant, 2012; Mon-Williams & McIntosh, 2000; Wing & Fraser, 1983) or both digits (Schot, Brenner, & Smeets, 2017; Smeets & Brenner, 1999; Smeets, van der Kooij, & Brenner, 2019) are transported to specific positions on the object. In the VHP condition, the positions on the object toward which the fingers were moving (grasping points) did not coincide with the haptic positions felt by the fingers of the left hand. Integrating these visual and haptic position estimates should have thus led to a worse grasping performance than when these estimates were congruent (VH vs. VHP), which was however not the case. It is rather more plausible that multisensory grasping movements are planned and executed based on a visuo-haptic estimate of the object’s centroid which is also used to better scale visual size. Alternatively, as suggested by one of the reviewers, the improvements observed in multisensory grasping movements might not be the result of a more reliable scaling of visual size, but rather the consequence of more successful sensory transformations that could facilitate the estimation of the grasping points on the object. Seeing the left hand holding the object and the haptic position cues provided by the same hand could assist the realignment of the grasping points that are initially available in different modality-specific coordinates (Kuling, Brenner, & Smeets, 2016; Kuling, van der Graaff, Brenner, & Smeets, 2017; Smeets, van den Dobbelsteen, de Grave, van Beers, & Brenner, 2006). This view might also explain why the contribution of haptic cues might, in specific cases, not be beneficial (Brenner et al., 1997).

In summary, our study provides a new perspective on the role of haptics in sensorimotor control by highlighting how non-visual inputs influence vision during object manipulation. Whether it be reaching for the cap of a handheld bottle, or passing a glass from one hand to the other, humans rely on haptics to guide movements more than we realize. Often, haptics can even take a leading role when vision is driven away from the manipulated object (e.g., when looking at a glass while uncorking a bottle). Hence, haptics is a remarkably underestimated source of information we draw from to guide efficient reaching and grasping movements in everyday life.

## Acknowledgements

We thank Prof. Jeroen Smeets for helpful comments and discussions.

## Data Availability

Experimental data described in this manuscript are available on Open Science Framework following this link: https://osf.io/apyks/. The code for the data analysis is also available at the same link.

## CRediT author statement

**Ivan Camponogara:** Conceptualization, Methodology, Software, Validation, Formal analysis, Investigation, Data Curation, Writing - Original Draft, Writing - Review & Editing, Visualization. **Robert Volcic:** Conceptualization, Methodology, Software, Validation, Formal analysis, Data Curation, Resources, Writing - Review & Editing, Visualization, Supervision.

## Additional information

**Competing interests** The authors declare no competing interests.

Correspondence and requests for materials should be addressed to I.C.

